# Inter-brain coupling reflects disciplinary differences in real-world classroom learning

**DOI:** 10.1101/2022.03.23.485430

**Authors:** Jingjing Chen, Penghao Qian, Xinqiao Gao, Baosong Li, Yu Zhang, Dan Zhang

**Author notes:** Corresponding authors: Dan Zhang, Yu Zhang.

## Abstract

Classroom is the primary site for learning. One important feature of classroom learning is its organization into different disciplines. While disciplinary differences could substantially influence students’ learning processes, little is known about the neural mechanism underlying successful disciplinary learning. In the present study, wearable EEG devices were used to record a group of high school students during their classes of a soft (Chinese) and a hard (Math) discipline throughout one semester. The students with higher learning outcomes in Chinese were found to have better inter-brain neural couplings with their excellent peers, whereas the students with higher Math outcomes were found to have better couplings with the class average. Moreover, the inter-brain couplings showed distinct dominant frequencies for the two disciplines. Our results illustrate disciplinary differences in successful learning from an inter-brain perspective and suggest the neural activities of excellent peers and class average as exemplars for soft and hard disciplines.

**Teaser:** Successful classroom learning is associated with distinct inter-brain coupling patterns for soft and hard disciplines

## Introduction

Classroom learning, where dozens of students learn together under the guidance of a teacher in a classroom, is the primary scenario for human beings’ formal learning. Due to its practical importance for personal development, classroom learning has drawn consistent attention from the fields of education and psychology^1–3^. It has also been considered an ideal starting point for real-world neuroscience In recent years for its widely-existence and semi-controlled structures^4^.

One important feature of classroom learning in educational practice is its organization into different disciplines (e.g., Math, history, physics, or language courses). It is widely acknowledged that disciplinary differences could substantially influence classroom learning. The hard-soft dimension is possibly one of the most influential frameworks regarding disciplinary differences^5^. Hard disciplines (e.g., math, natural science, and engineering) are known for the relatively hierarchical, linear knowledge structure and straightforward, uncontentious learning contents. Soft disciplines are usually associated with loose-structured, non-linear knowledge and contents that require more constructive and interpretative activity (e.g., history, philosophy, and language courses)^5–7^. The differences in the disciplinary knowledge have been proven to influence teachers’ teaching goals and correspondingly shape students’ learning processes towards success^8,9^. For instance, it has been proposed that students prioritize fixed knowledge from teachers when learning hard disciplines over their soft counterparts^10^. Nevertheless, it should be noted that disciplinary differences in the successful learning process have mainly been inferred based on indirect data such as expert evaluation, retrospective self-reports, and learning outcomes.^7,10,11^. There is a dearth of empirical studies directly addressing the learning process itself^12^.

The emerging inter-brain coupling analysis has demonstrated its potential as a powerful tool to directly capture the learning process. The inter-brain coupling approach identifies neural correlates of interests by computing one’s inter-brain coupling to other people who are situated in the same learning environment or share the same learning tasks (e.g., attending lectures or videos in a classroom)^4,13–16^. Recent studies have reported that the average inter-brain coupling from one student to all their peers or classmates (termed as ‘student-class coupling’ in the following) during the learning process was positively correlated with students’ engagement^4^, memory retention performance^15^, and their final-term exam scores^14^. These findings suggest that student-class coupling is capable of characterizing one’s moment-to-moment learning process^4,14^. At the same time, it should be noted that in these studies, the disciplines investigated are hard ones, such as biology, computer science, and physics^4,13–16^. As the learning of hard disciplines is straightforward and non-ambiguous (e.g., the application of fixed mathematical rules)^17^, it is plausible to assume that similar neural activities responding to the learning contents could emerge among classmates during the learning process. Hereby, averaging might facilitate the extraction of the neural activity shared across students that could effectively represent the ‘desired’ learning process as intended by the courses. Therefore, it seemed promising to consider the ‘class average’ neural activity for representing a successful learning process in hard disciplines. However, since creative ideation and personalized construction are cherished in soft disciplines, the differences in teaching goals might lead to a different learning process for students to meet teachers’ requirements^7^. Accordingly, the possibly different learning processes towards success might undermine the importance of ‘class average’ in the context of soft disciplines. Nevertheless, no single inter-brain study has addressed the issue of disciplinary differences.

Excellent peers could serve as a candidate exemplar to represent a successful learning process in soft disciplines. Here, excellent peers (termed as ‘excellence’ in the following) refer to the students in a class with top learning outcomes. As good learning outcomes have long been associated with an effective learning process^18^, it is reasonable to take the neural activities of excellence to represent the learning process towards success. Compared to the ‘class average’, the excellence could be a better candidate for a successful learning process of soft disciplines, given a potentially different learning goal. While comparing these two representatives (i.e., ‘class average’ vs. excellence) for successful learning could be challenging with conventional single-brain-based methods, the inter-brain approach enables us to compare them directly. Specifically, it is straightforward to compute student-class coupling and student-excellence coupling similarly. Comparing the student-class coupling and student-excellence coupling during the learning of soft and hard disciplines may give us insights into disciplinary differences in successful learning.

The real-world classroom is expected to serve as an ideal site to investigate the disciplinary differences in successful learning. Compared to the conventional laboratory-based studies that have mainly focused on strictly-controlled, parametric experimental designs (i.e., based on contrasts across simplified learning tasks to isolate targeted factors in disciplinary differences) to remotely resemble real-world learning^19– 21^, classroom-based studies are advantageous for their high ecological validity since they could directly reflect the complex and dynamic disciplinary learning process that happens every day. The recent development of wearable electroencephalogram (EEG) devices has enabled researchers to track students’ learning processes in real-classroom settings^22–25^. For instance, the EEG device in the form of a headband could support the easy acquisition of EEG data from an entire class of students for its portability, usability, low purchase, and running cost. Wearable EEG devices have been proved to be effective in tracking students’ sustained attention, situational interests, and engagement during their classroom learning processes^4,26,27^. The ecologically naturalistic paradigm with wearable neuroimaging technologies is expected to provide insights into understanding the disciplinary differences ‘in the wild’ and offer a possible ‘fast lane’ to apply neuroscience findings into educational theoretical construction and practical application^23,28^.

Therefore, the present study aimed to investigate disciplinary differences in the successful learning process by recording EEGs from a group of high-school students in China during their regular classroom courses for a whole semester. Math and Chinese (the mother tongue learning in China) were chosen as representative courses for the hard discipline and soft discipline, respectively, as they are two of the most important compulsory courses before college in China. Based on the state-of-the-art understanding of disciplinary learning^4,14,15^, we hypothesized that the ‘class average’ might effectively represent the successful learning process during Math courses (a hard discipline). In contrast, excellent peers could effectively represent the successful learning process of Chinese (a soft discipline). Correspondingly, the student-class coupling was expected to be correlated with the learning outcome of Math courses, and the student-excellence coupling was expected to be correlated with the learning outcome of Chinese courses. No clear hypothesis regarding the specifically involved frequency band was formulated due to limited evidence. Any discovery would promote our understanding of the neural mechanism behind the successful disciplinary learning process in the classroom.

## Result

### Real-world classroom setting-up and inter-brain analysis

As demonstrated in Fig.1**a**, thirty-four students from the same class (thirty-seven students in total) from grade 10 of a high school in Beijing volunteered to join the study. Wearable EEG headbands with two dry electrodes covering Fp1/2 were chosen to record students’ brain signals during their regular Math and Chinese sessions in the classroom throughout one semester. While the EEG headbands have demonstrated their effectiveness in tracking brain states in tasks such as resting-state, sudoku games, and surgery^29–33^, their signal quality was also validated here with a 2-minute eye-closed/open task. Twenty-two students out of the same class participated in this task in their classroom environment. Supplementary Fig.S1 shows the signal quality validation results: compared to the eye-open condition, an expected spectral peak in the alpha range (8-13 Hz) was observed in the eye-closed condition, demonstrating the reliability of the headband for EEG recordings in the classroom.

**Fig.1:**
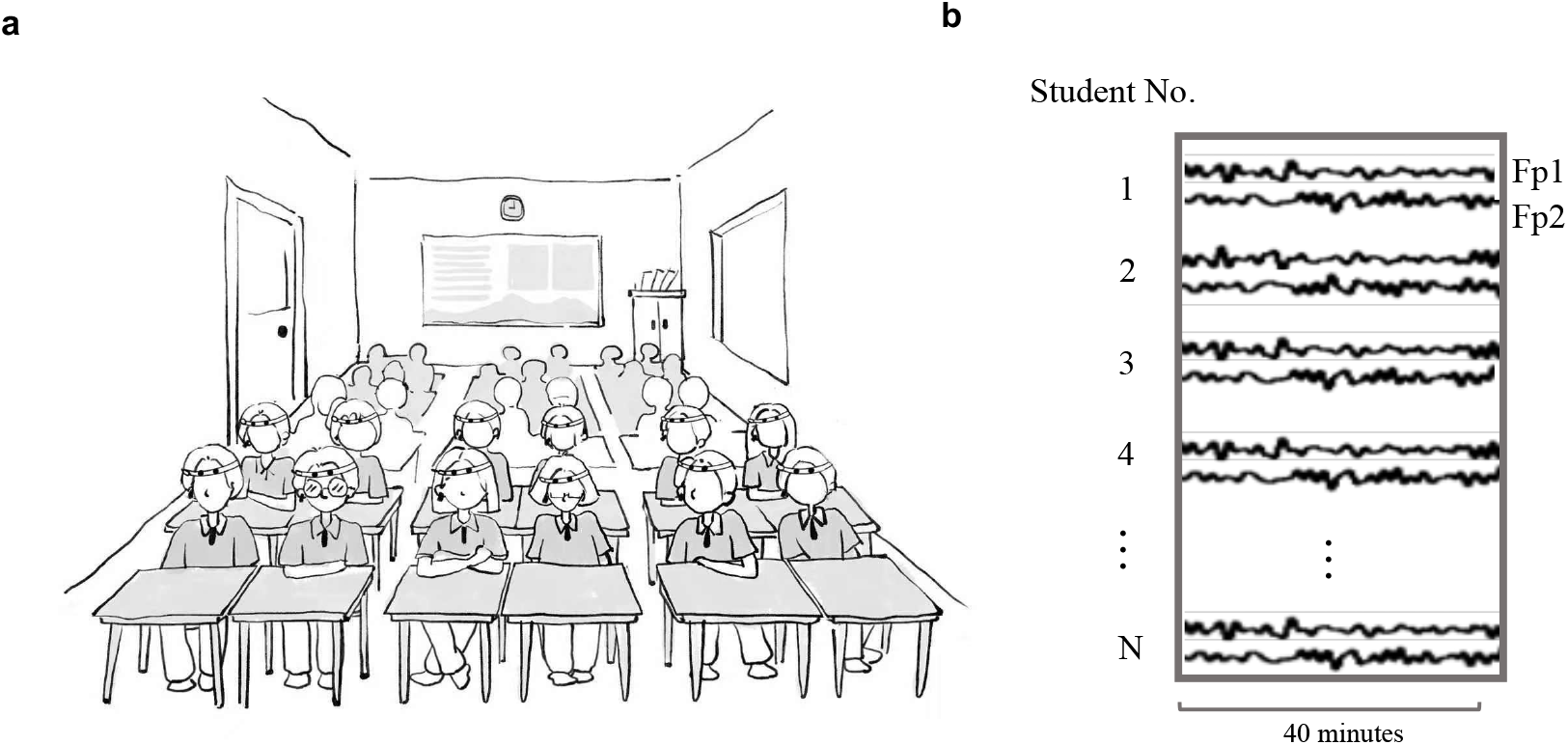
Experiment paradigm. **a**, An illustration of the experimental setup for students wearing an EEG headband during their regular classroom learning; **b**, An illustration of the recorded EEG signal during a session; EEGs were recorded at Fp1 and Fp2 for all the students for 40 minutes during a session.

**Fig 2:**
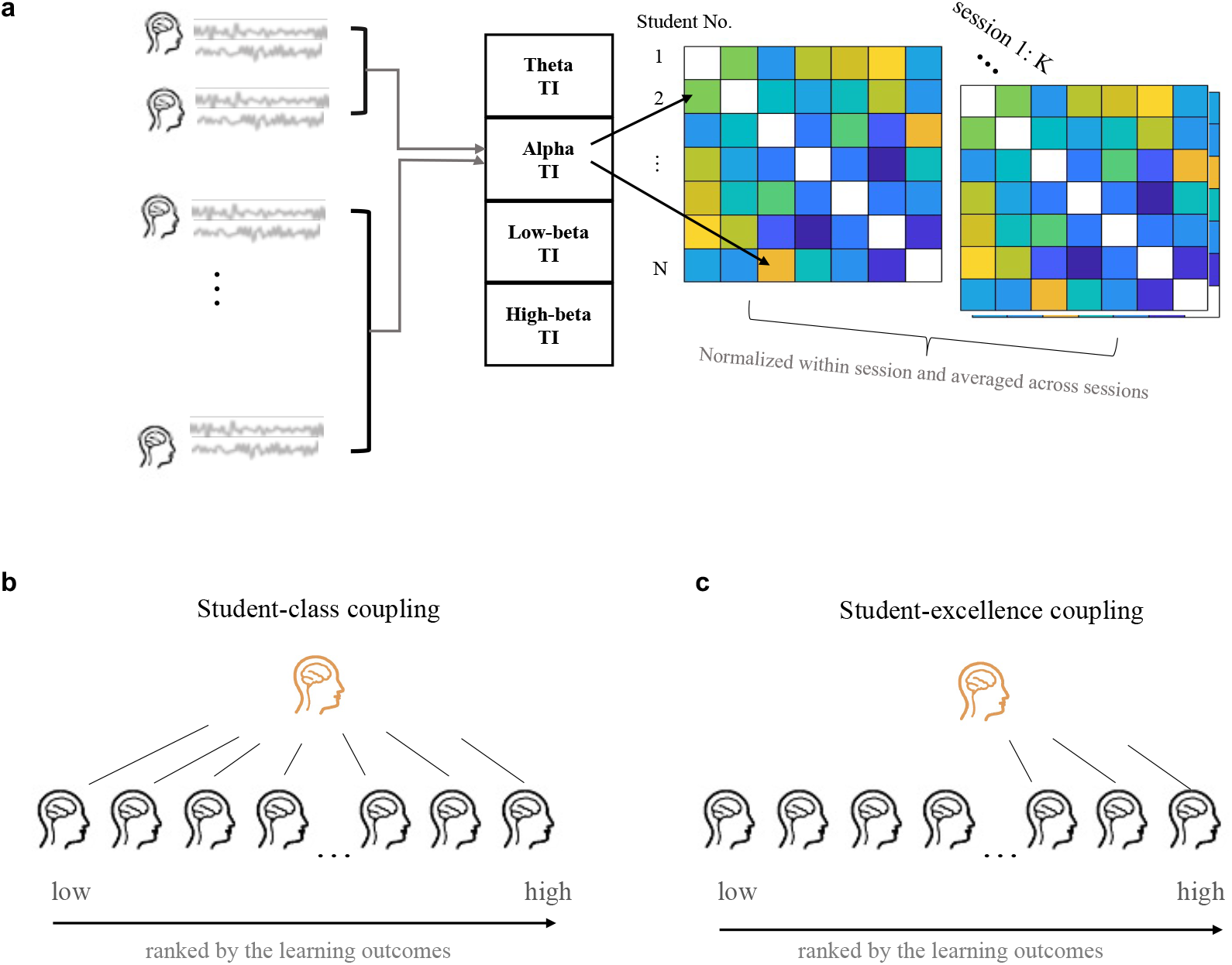
A schematic illustration of the inter-brain coupling analysis. **a**, Computations of pairwise total interdependence (TI) matrix for each pair of students (i, j) for each session, at the frequency bands of theta, alpha, low-beta, and high-beta. The TI values were then normalized within each session and averaged across sessions to obtain an inter-brain coupling value for each pair of students. **b**, Student-class coupling was obtained by averaging TI values over all possible pairwise combinations between one student and the rest of the class. **c**, Student-excellence coupling was computed by averaging TI values over all possible pairwise combinations between one student and all the excellences in the class, except for this student himself/herself if the student was one of the excellences.

The data collection lasted for four months to cover the whole semester. Each month, students’ EEG signals were acquired during Chinese and Math sessions for one week (one or two sessions per day) following the regular curriculum. The total number of sessions was 38, with 18 sessions for Chinese and 20 sessions for Math. The Math course included the introduction of planar vectors, cubic geometry, plural, statistics, and probability. The Chinese courses included reading ancient and modern poems, essays, and novels, and an introduction to writing. Each session lasted for 40 minutes. The students’ final-term exam scores for Chinese and Math at the end of this semester were taken to indicate their learning outcomes. Compared with the specially-designed quiz, final-term exams were expected to boost the ecological validity as they were derived from the highly-developed evaluation system in the daily educational practice. The final exams covered the contents of the whole semester.

The total interdependence (TI) method has been employed in the present study to calculate the inter-brain coupling by computing the magnitude squared coherence between brain signals simultaneously recorded from two students^4,16,34^. Recent inter-brain studies have validated the efficiency of TI methods in tracking individuals’ engagement and valence levels in naturalistic scenarios such as a classroom and a concert hall^4,35^. Then, to test our hypothesis, student-class coupling and student-excellence coupling were calculated for each student as indicators of the disciplinary learning process for the whole semester, as shown in Fig.3**b, c**. Then, Pearson’s correlations between individuals’ student-class coupling (or student-excellence coupling) and their corresponding final exam scores were calculated separately for soft and hard disciplines to identify neural correlates of successful learning. For student-class coupling, all students were included in the correlation analysis. However, for student-excellence coupling, the excellences themselves were excluded during the correlation analysis between student-excellence coupling and final exam scores. For example, if individuals’ student-excellence coupling was computed with the top *N*_*E*_ students, the top *N*_*E*_ students’ coupling values, as well as their final exam scores, would be removed. The number of students used to conduct the correlation analysis would then be the number of students left after subtracting the top *N*_*E*_ students from the total number of *N* students (*N* − *N*_*E*_). Theta (4-8 Hz), alpha (8-13 Hz), low-beta (13-18 Hz), and high-beta (18-30 Hz) bands were calculated separately in the inter-brain coupling analysis.

**Fig.3:**
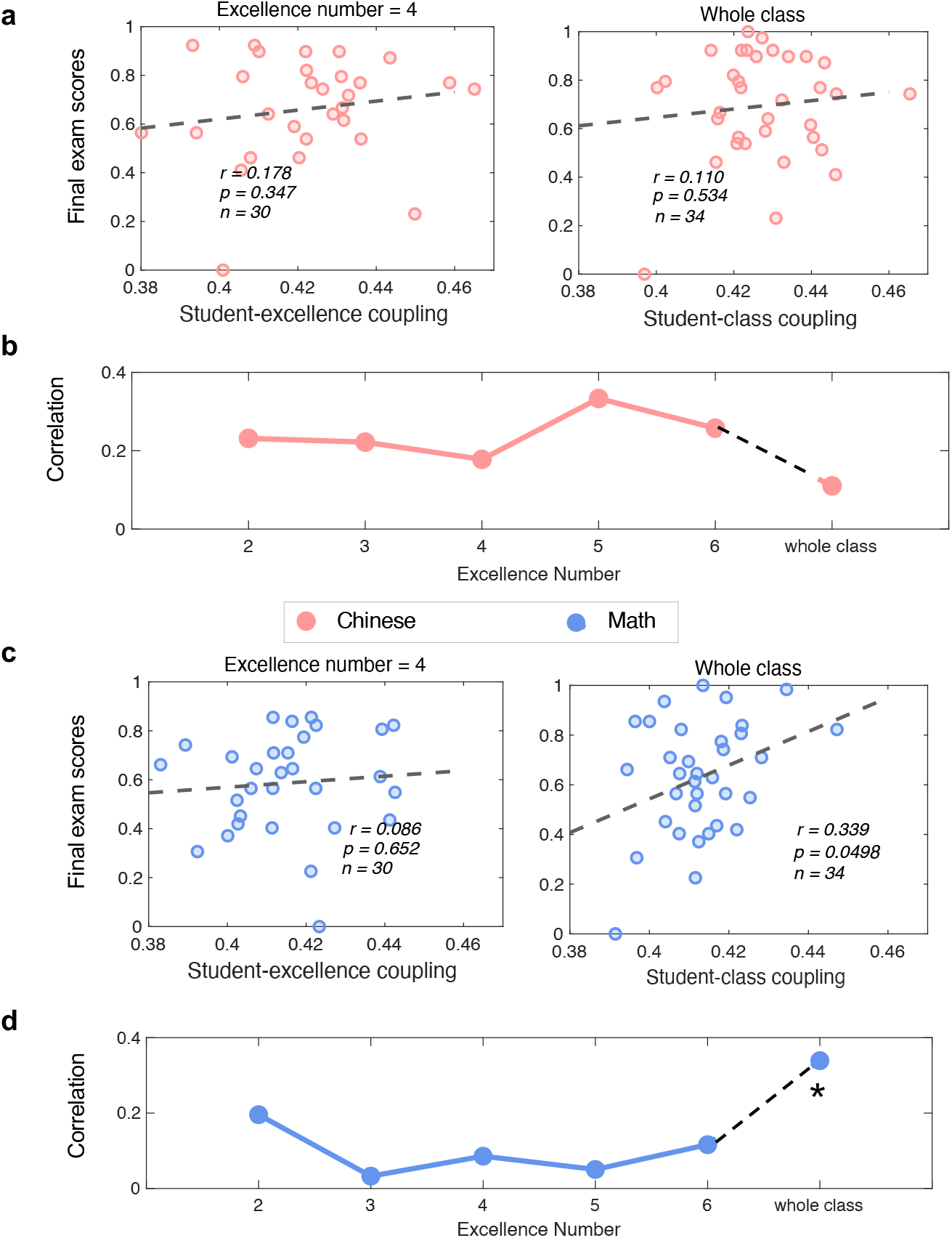
Correlations between theta-band student-excellence/class coupling with one’s final exam score for Chinese and Math. **a**, Scatter plots between theta-band student-excellence coupling (left, excellence number = 4) and theta-band student-class coupling (right) and the final exam scores of Math. **b**, Correlation r values as a function of the number of excellences included in the calculation of student-excellence coupling. **c**, Scatter plots between theta-band student-excellence coupling (left, excellence number = 4) and theta-band student-class coupling (right) and the final exam scores of Chinese. **d**, Correlation r values as a function of the number of excellences included in the calculation of student-excellence coupling. Note that the excellences themselves were not included in the correlation analysis. The star indicates a significant (*p* < 0.05) correlation.

### Theta-band student-class coupling reflects successful classroom learning for Math

Here, no significant correlations were observed between theta-band inter-brain coupling and the final exam scores for Chinese, neither in student-excellence coupling nor in student-class coupling (Fig.3**a, b**; student-excellence coupling: *r* = 0.178, *p* = 0.347, *n* = 30, excellence number = 4; student-class coupling: *r* = 0.110, *p* = 0.534, *n* = 34). On the other hand, theta-band student-class coupling during Math sessions was found to be positively correlated with the final exam scores for Math (Fig.3**c, d**): the students with higher learning outcomes in Math were found to have better inter-brain couplings with other classmates (*r* = 0.339, *p* = 0.0498, *n* = 34). No significant correlations between theta-band student-excellence coupling and Math scores were found, with the number of excellences included in the calculation of student-excellence coupling varying from 2 up to 6.

### Alpha-band student-excellence coupling reflects successful classroom learning for Chinese

Alpha-band student-excellence coupling during Chinese sessions was significantly correlated with the final exam scores for Chinese (Fig.4**a**): the students with higher learning outcomes in Chinese were found to have better inter-brain couplings with their excellent peers (*r* = 0.433, *p* = 0.017, *n* = 30, excellence number = 4). The correlation remained significant or marginally significant when the number of excellences included in the calculation of student-excellence coupling varied from 2 to 6 (Fig.4**b**). No significant correlations between alpha-band student-class coupling and the final exam scores of Chinese were found (*r* = 0.077, *p* = 0.665, *n* = 34). Moreover, no significant correlations were observed between alpha-band inter-brain coupling and the final exam scores of Math, neither in student-excellence coupling nor in student-class coupling (Fig.4**c, d**; student-excellence coupling: *r* = 0.266, *p* = 0.156, *n* = 30, excellence number =4; student-class coupling: *r* = 0.195, *p* = 0.270, *n* = 34).

**Fig.4:**
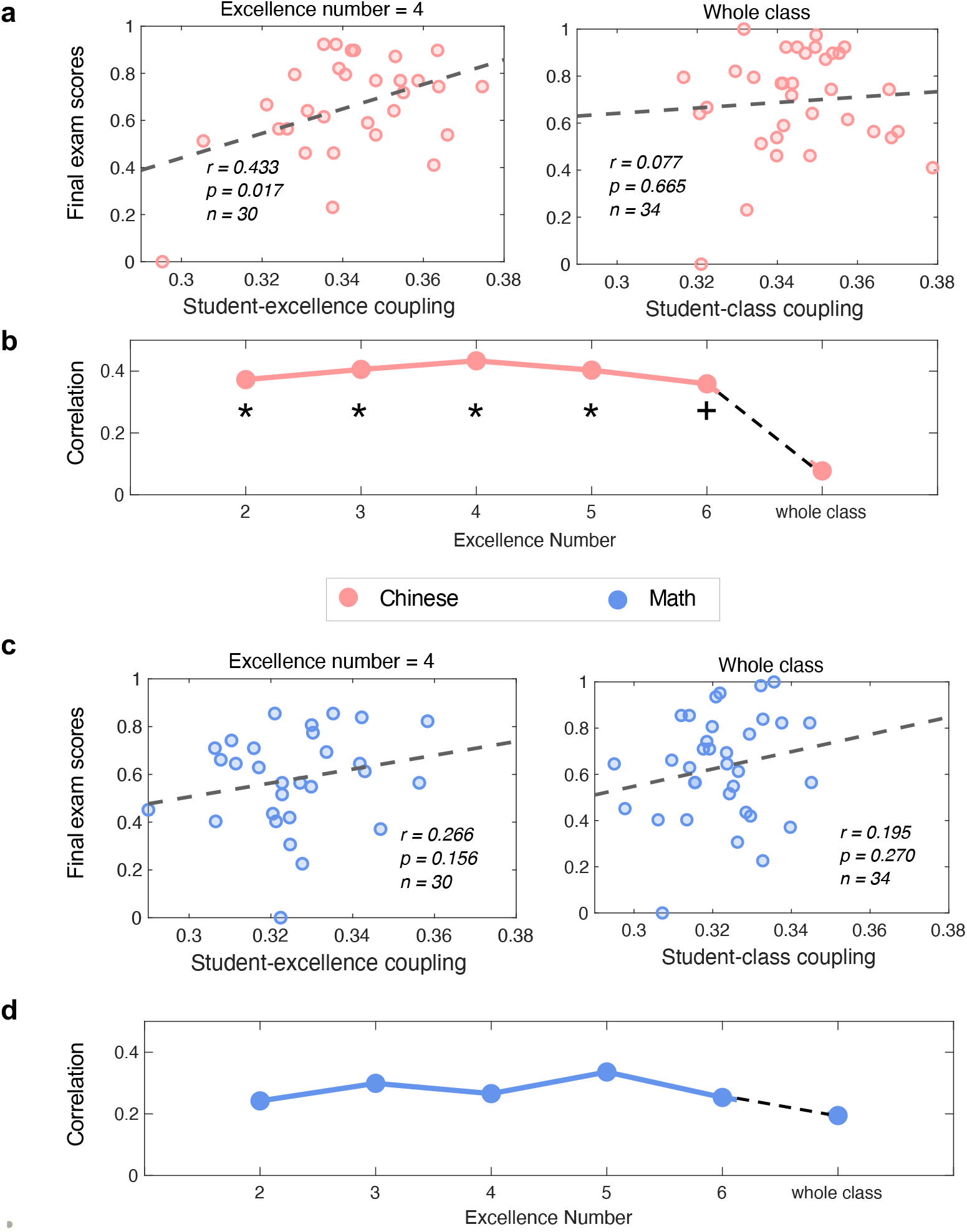
Correlations between alpha-band student-excellence/class coupling with one’s final exam score of Chinese and Math. **a**, Scatter plots between alpha-band student-excellence coupling (left, excellence number = 4) and alpha-band student-class coupling (right) and the final exam score of Chinese. **b**, Correlation r values as a function of the number of excellences included in the calculation of student-excellence coupling. **c**, Scatter plots between alpha-band student-excellence coupling (left, excellence number = 4) and alpha-band student-class coupling (right) and the final exam scores of Math. **d**, Correlation r values as a function of the number of excellences included in the calculation of student-excellence coupling. Note that the excellences themselves were not included in the correlation analysis. Stars indicate significant (*p* < 0.05) correlation and the cross indicates a marginal significant (*p* < 0.10) correlation.

### Frequency-specificity of outcome-related inter-brain coupling

Fig.5 further showed the overall inter-brain coupling results in four frequency bands (theta, alpha, low-beta, and high-beta). Inter-brain coupling in the theta and alpha bands was found to correlate with the final exam scores (as shown above) significantly. In contrast, the inter-brain coupling at the low-beta and high-beta bands failed to reach significance. We also conducted a similar inter-brain coupling analysis but focused on 1-20 Hz as several previous studies^4,16^. The results were shown in Supplementary Fig.S2: No significant correlations were found between 1-20 Hz inter-brain coupling and the final exam scores, demonstrating the value of separating different frequency bands.

**Fig.5:**
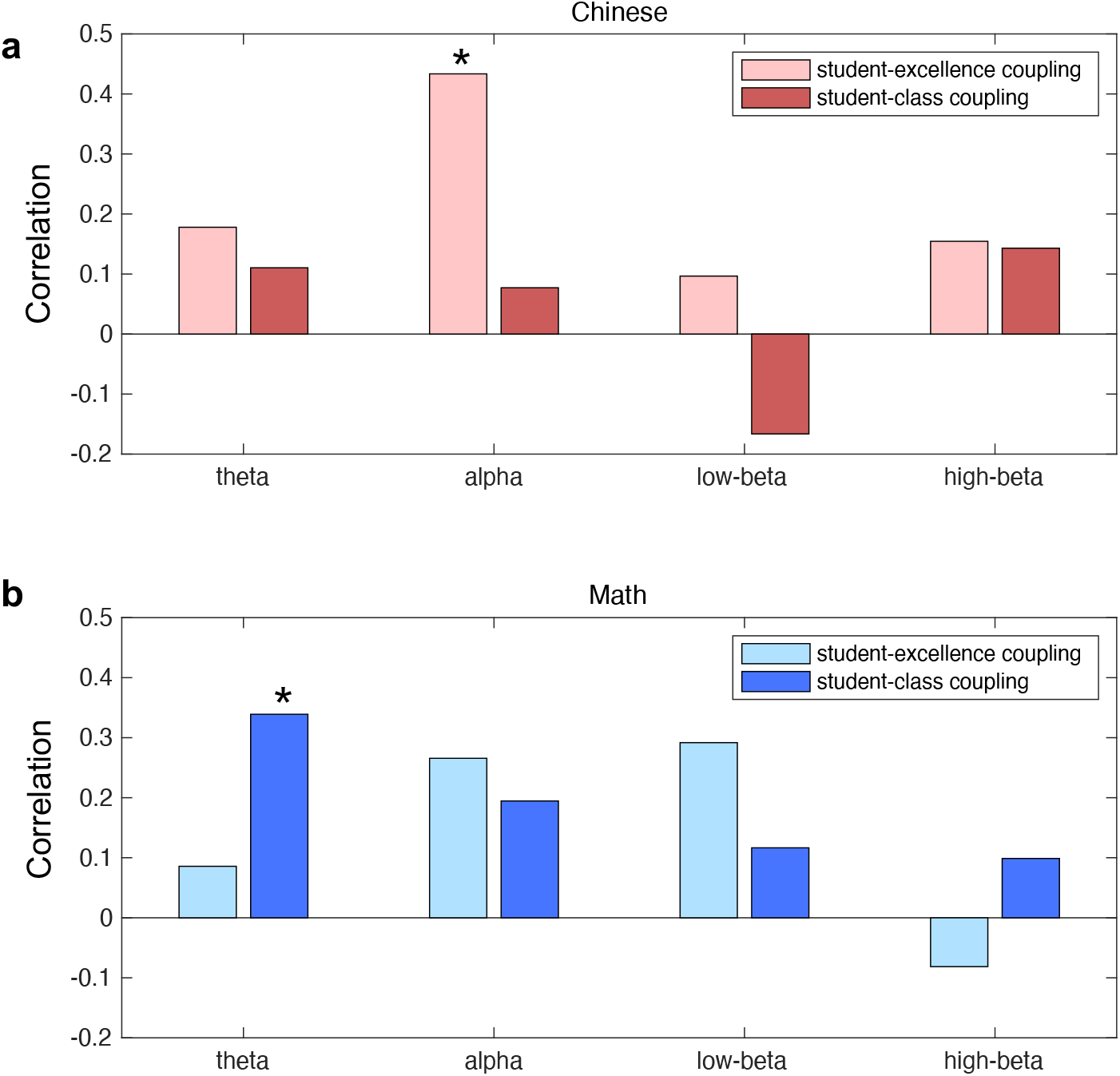
Correlation r values between inter-brain coupling and the final exam scores at the theta, alpha, low-beta, and high-beta bands for **a**, Chinese and **b**, Math. Bars with a lighter color indicated student-excellence-coupling-based correlations (excellence number = 4), and bars with a darker color indicated student-class-coupling-based correlations. Stars indicated a significant (*p* < 0.05) correlations.

To explore the discipline-specificity of the inter-brain coupling results, the correlations between the final exam scores and inter-brain coupling were re-computed by switching the disciplinary scores (i.e., computing the correlation between inter-brain coupling during Math sessions and the final exam scores of Chinese and vice versa). As shown in Fig. 6**b**, theta-band student-class coupling during Chinese sessions is significantly correlated with the final exam scores of Math (student-class coupling: *r* = 0.345, *p* = 0.045, *n* = 34). No other correlations reached a significant level after switching.

**Fig.6:**
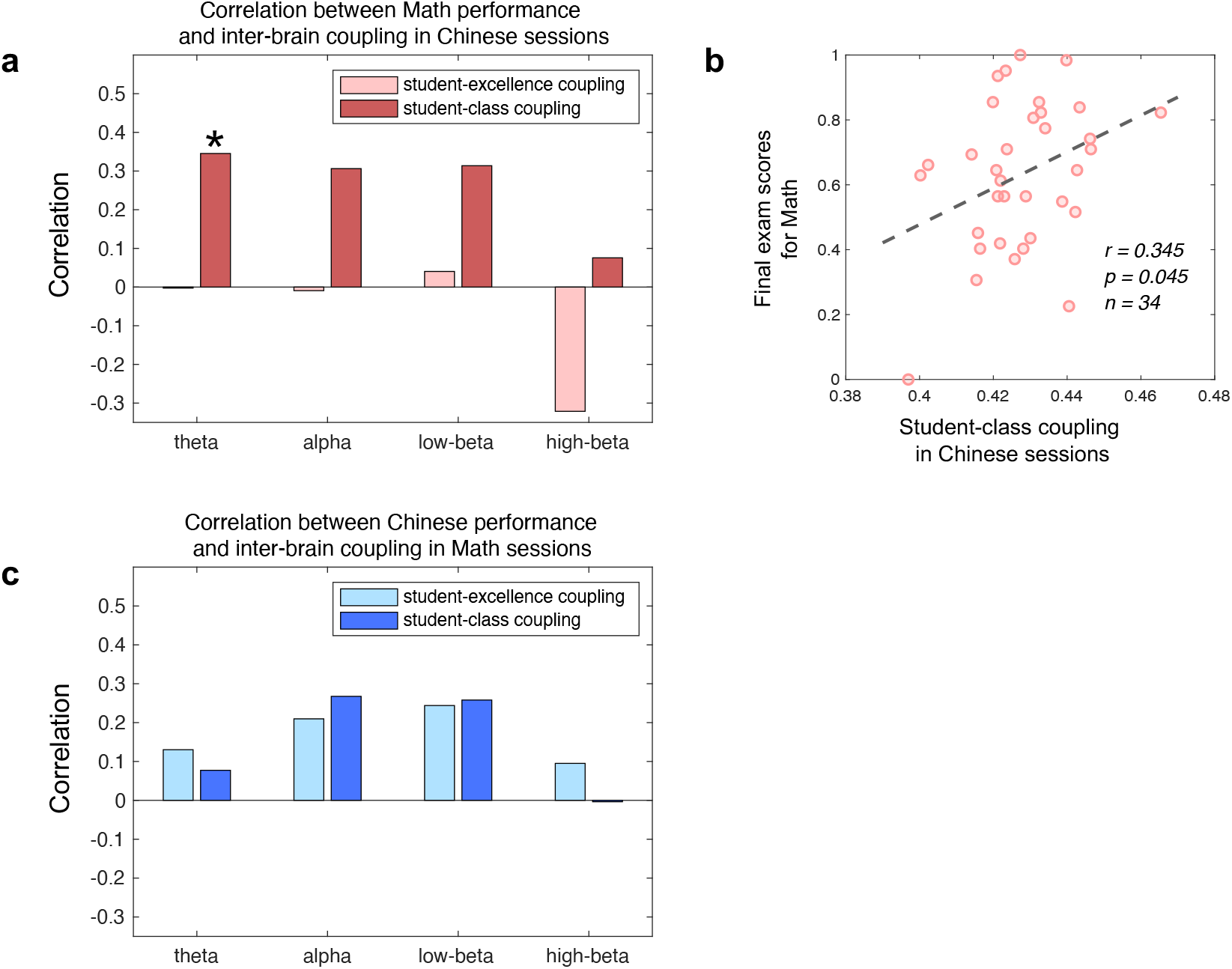
Results of correlation analysis by switching the discipline scores, i.e., correlating inter-brain coupling in Math sessions with the final exam scores of Chinese (**a**) and vice versa (**c**). **b**, Scatter plots between theta-band student-class coupling during Chinese sessions and the final exam scores of Math. Bars with a lighter color indicated student-excellence-coupling-based correlations (excellence number = 4), and bars with a darker color indicated student-class-coupling-based correlations. The stars indicated a significant (*p* < 0.05) correlation.

### Single-brain features fail to reflect successful learning

Additionally, we conducted similar correlational analyses between single-brain EEG features and the final exam scores. The relative power of the four frequency bands from each student was taken as the single-brain EEG features. As shown in Fig.7, the single-brain analysis reveals no significant correlations with the disciplinary final exam scores (Fig.7**b**; the highest correlation for Chinese in the theta band: *r* = -0.180, *p* = 0.307, *n* = 34; Fig.7**d**; the highest correlation for Math in the high-beta band: *r* = -0.141, *p* = 0.425, *n* = 34).

**Fig.7:**
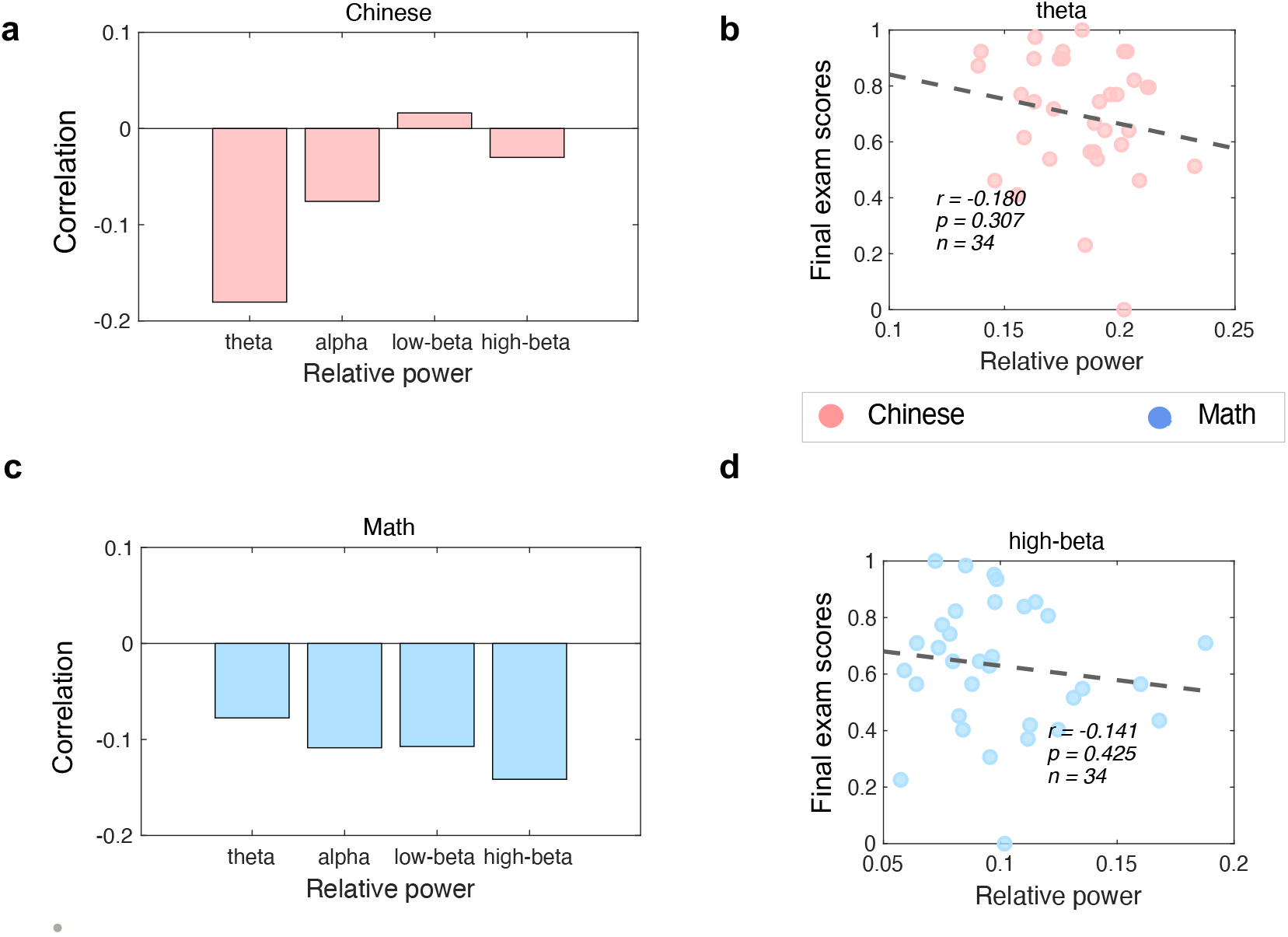
The correlation between the single brain’s relative frequency power in the theta, alpha, low-beta, and high beta band with one’s final exam scores of Chinese (**a**) and Math (**c**). No significance was found in any of the frequency bands. **b**, Scatter plots between relative power in the theta band and the final exam scores of Chinese (the highest correlation for Chinese). **d**, Scatter plots between the relative power in the high-beta band and the final exam scores of Math (the highest correlation for Math).

## Discussion

In the present study, the learning processes of high school students while taking a soft (Chinese) and a hard (Math) discipline in their real classroom were recorded by wearable EEG devices for a whole semester. By taking their final-term exam scores as learning outcomes, students with higher Chinese outcomes were found to be associated with better inter-brain neural couplings with their excellent peers during the Chinese courses, whereas students with higher Math outcomes were found to be associated with better couplings with other classmates during both the Chinese and the Math courses. Moreover, the neural couplings showed different dominant frequencies for the two disciplines. While the outcome-related inter-brain coupling for Math was found in the theta-band, the importance of the alpha-band was highlighted in the successful learning for Chinese courses. No significant correlation was found between the single brain’s relative power and final exam scores in either discipline. Our results demonstrate the feasibility of inter-brain coupling to eval students’ successful learning process for both soft and hard disciplines. More importantly, the present study provides insights into understanding the disciplinary differences ‘in the wild’ from an inter-brain perspective, suggesting the neural activities of excellent peers and class average as exemplars for successful classroom learning in soft and hard disciplines, respectively.

The correlation between individuals’ student-class coupling and their learning outcomes for Math verified and extended previous findings of neural mechanisms underlying the learning process. By investigating hard disciplines such as physics, biology, and computer science, recent studies have demonstrated student-class coupling as a useful tool to evaluate the learning process^13–15^. Our results about Math, another representative hard discipline, are in line with these studies, where student-class coupling was also found to be correlated with students’ learning outcomes. Moreover, after decomposing data into different frequency bands, our results extended previous findings by showing the importance of the frontal theta-band activity during real-classroom learning. Frontal theta activity has been reported to reflect cognitive processes such as cognitive control^36–38^, sustained attention^39^, and working memory^40– 42^, and has been found to increase in the arithmetic-related tasks^43^. During the learning of hard disciplines, the emphasis on the development of a capacity to master and apply the accepted scientific viewpoints would require the students to align with the course material^6^. Hereby, the theta-band brain activity shared across classmates could reflect students’ continuous engagement with the course content. Then, theta-band student-class coupling could imply the extent to which each student attended the course content^4,15^, or the extent to which each student interpreted the course content^14^. Therefore, better learning outcomes for Math are found to be associated with better inter-brain coupling with other classmates in the theta band.

The positive correlation between theta-band student-class coupling during the Chinese sessions and the learning outcomes of Math further suggested that the class-level frontal theta-band activity could reflect the required cognitive processes shared across the two disciplines. The learning process of Chinese would also rely on cognitive control, sustained attention, and working memory, which allowed students to attend the courses. Nevertheless, the non-significant correlation between theta-band student-class coupling and the Chinese final exam scores in the present study might also suggest the relatively loose link between these cognitive processes and the learning outcomes of Chinese. This finding is similar to a previous study where students’ studiousness and continuous engagement have been suggested to be beneficial for both the learning of Math and German. However, they are more important for the learning outcomes of Math^44^.

The positive correlation between alpha-band student-excellence coupling during the Chinese sessions and the students’ Chinese final exam scores provides evidence of the critical neural correlates of successful learning in soft disciplines. The distinct frequency band (alpha) compared to Math (theta) suggests that successful learning of Chinese and Math relies on substantially different cognitive processes. Despite the lack of neuroscience evidence on soft-discipline learning, the frontal alpha-band activity could be related to the inhibition of stimulus-driven attention^39,45^ and was found to be involved in tasks with high internal processing demands such as creative ideation^46,47^ and imagination^48^. At the same time, student-excellence coupling rather than student-class coupling was informative about the learning outcomes in Chinese, highlighting the neural activities of excellent peers as exemplars for successful learning in soft disciplines. Although different excellence might have different internal interpretations of the course contents, they could share the similar temporal dynamics of the interpretation process. For instance, while learning an ancient poem, two top students could immerse in the aesthetic experience simultaneously when imagining different scenarios in their minds. Note that EEG recording techniques used in our study are advantageous for capturing the temporal dynamics of the learning process rather than the fine-grained representation of the learning content. Taken together, it is plausible to assume that the temporal dynamics of the frontal alpha activity shared across excellences might represent an internal processing state for interpretation construction, which is critical for the learning of Chinese. Moreover, unlike the responses to external stimuli (course contents), this internal state may not necessarily share across classmates, which results in positive results of student-excellence coupling rather than student-class coupling.

It is necessary to distinguish between the excellent peers found in this study and the experts that have been often referred to in the field of education. In educational practice, the expert-like mastery of knowledge has been regarded as the target of students’ learning and has been linked to good learning outcomes^49^. For example, the performances of emergency medicine trainees were found to improve in the crisis resource management tasks when their cognitive processes were more expert-like^50^. A recent functional magnetic resonance imaging (fMRI) study also reported that one’s neural alignment (coupling) to the experts could positively predict the final-term exam score of their computer science course^14^. While experts have been regarded as a well-established exemplar for successful learning, our results suggest that excellences also serve as an alternative reference in soft disciplines. Moreover, compared with experts who may learn qualitatively different from students due to their broader understanding of the field^14^, excellences with similar prior knowledge about the to-be-learned content may be particularly efficient as an exemplar for the learning process starting as a novice.

It should also be noted that the results of non-significant student-excellence-coupling-based correlation for the learning outcomes of Math did not necessarily undermine the potential importance of excellence for successful Math learning. On the one hand, the correlation coefficients between the Math-session student-excellence coupling and the Math final exam scores still reached a positive value of >0.2 at the alpha band (Fig. 4**d**). On the other hand, the students of Math excellence might not adequately express their optimal learning processes during the Math sessions. Specifically, since the classroom teaching was designed to meet the need of the majority of the class^51–53^, there might be a lack of challenge for Math excellence. Consequently, the possible boredom might demotivate the excellence to follow the lectures^54^ and eliminate the possibly-existing correlations. By contrast, the teaching of soft disciplines such as Chinese emphasizes constructive and interpretative activity, which is expected to be similarly challenging for students at different proficiency levels.

As the first study investigating the disciplinary differences in students’ successful learning process in real-classroom settings, several limitations must be noted. First, the present ecologically valid paradigm posed a challenge to strictly-controlled contrasts between disciplines. Multiple factors (e.g., learning contents, learning goals and learning difficulties) could lead to the distinct outcome-related inter-brain coupling patterns in soft and hard disciplines. While this is how disciplinary differences manifest in everyday learning processes, future work will be needed to clarify the unique contributions of these factors. Second, the present effect sizes are relatively small compared with previous inter-brain studies (the correlation between inter-brain coupling and learning outcomes for Chinese: R-squared = 0.187; for Math: R-squared = 0.115). In a recent fMRI study focused on the neural correlates of successful learning, R-squared values were found to be influenced by the brain regions, varying from 0.168 to 0.563 across brain regions^14^. While the dual-channel EEG devices in the present study provide convenience for daily longitudinal acquisition, the relatively few channels might fail to fully capture the learning-related brain activities. For instance, the development of portable EEGs with larger coverage areas^4^ and portable functional near-infrared spectroscopy (fNIRS) devices^55^ could be a plausible solution in the near future. Third, while Chinese and Math have been chosen to represent soft and hard disciplines here, more disciplines were needed to be investigated to verify the framework of hard/soft disciplines thoroughly. Specifically, as our findings indicated that some cognitive processes might share across the learning of different disciplines, the involvement of more disciplines will facilitate a deeper understanding of the domain-specific process and domain-general process across disciplines during real-world learning activity^56^.

## Methods

### Participants

Thirty-six students (16 females; age: 15-16 years old) from the same class (37 students in total) in grade 10 of a high school in Beijing volunteered to wear a headband EEG device during their regular Math and Chinese sessions throughout one semester. The study was conducted in accordance with the Declaration of Helsinki, and the protocol was approved by the ethics committee of the Department of Psychology, Tsinghua University (THU201708). All the participants and their legal guardians gave their written informed consent.

### Procedure and Data Recording

In the present study, a dual-channel headband with dry electrodes was used to record EEG at Fp1 and Fp2 over the forehead at a sampling rate of 250 Hz (Brainno, SOSO H&C, Korea). The reference electrode was placed on the right ear lobe with a ground at Fpz. The EEG device has been used previously in monitoring brain state during resting state, sudoku games, and surgery^29–33^. The signal quality of the headband was also tested in the present study by using an eye-closed/open task with 22 out of the same 36 students in their classroom environment.

The data collection lasted for four months to cover the whole semester. For each month, students’ EEG signals during Chinese sessions and Math sessions were recorded for one week (one session or two sessions per day) following the regular curriculum. The total number of sessions was 39, with 19 sessions for Chinese and 20 sessions for Math. Before Chinese or Math sessions began, students wore headbands with the help of experimenters, and the headbands were taken off after each session. Each session lasted for 40 minutes. There was one Chinese session when EEG devices failed to record any data due to technical issues. Two students were omitted from the analysis due to the consistently poor quality of EEG data across sessions. A total of 34 students in 18 sessions for Chinese and 20 sessions for Math were included in the following analysis.

During the Chinese and Math sessions, the learning content was taught according to the arrangement of the school. The Math sessions include the introduction of planar vectors, cubic geometry, plural, statistics, and probability; the Chinese sessions include reading ancient and modern poems, essays, and novels and the introduction to writing.

The students’ final exam scores in Chinese and Math were taken as indicators of their learning outcomes. The final exams covered the contents of the whole semester. Both exams were scored out of 100. The median of the students’ Math scores was 73, ranging from 33 to 95, and the median of the students’ Chinese scores was 69, ranging from 40 to 79. The scores were sufficiently diverse to characterize students’ differences in learning outcomes. These scores were normalized to [0, 1] using a min-max transformation for the following analysis.

### Data Preprocessing

Since EEG data were recorded in a regular classroom environment and students were instructed to attend Chinese and Math sessions as usual, more artifacts were expected as compared to conventional, highly-controlled laboratory settings. In the present study, there were three types of prominent artifacts: 1) a high value indicating signal saturation possibly due to losing contact with the headband; 2) slow drifts related to extensive head or body movements; 3) ocular artifacts related to eye movements.

The recorded EEG data were segmented into non-overlapping 30-sec epochs for preprocessing. As shown in Supplementary Fig. S3, ratios for saturated samples per epoch illustrated a 2-tailed distribution that most epochs containing saturated samples for less than 10% or more than 90%. Therefore, 50% was chosen as a threshold empirically. One epoch would be rejected if it contained saturated samples for more than 50%. The remaining epochs were then processed to remove the slow drifts with the NoiseTools toolbox ^57^ under Matlab (MathWorks, USA). The removal of the slow drifts was achieved by using the nt_detrending() function. By estimating the position of the glitch, this function could perform a weighted polynomial fit and achieve a better fit to the non-glitch parts. The processed epochs were further band-pass filtered between 0.1 Hz and 40 Hz with 1-s zero-padding at both ends. Afterward, the ocular artifacts were attenuated with the MSDL (multi-scale dictionary learning) toolbox, which was efficient in ocular artifacts removal for single-channel EEG signals^58^. Epochs were decomposed into neuronal and non-neuronal sources with dictionary learning. Then, the coefficients of non-neuronal sources were set to zero to achieve artifact reduction with the seq_MSDL() function. Supplementary Fig.S4 and S5 illustrated representative examples before and after the artifacts rejection procedure. Finally, epochs were rejected automatically if any samples in any channels exceeded a ±150 μV threshold. With the above preprocessing procedure, 57.2±1.85 % epochs were retained per student, ranging from 31.2% to 76.7%. The data retention rate was comparable with previous EEG studies in classroom settings^4,16^. The number of retained epochs per session per student was shown in Supplementary Fig.S6.

### Data Processing

Inter-brain coupling between all possible student pairs was computed using the total interdependence (TI) method as its efficacy in capturing the temporal relationship between two time series^34^. TI was estimated by computing the magnitude squared coherence using the Welch method when clean 30-s epochs were available at the same moments from both students. For *X*_*i*_, a 30-s epoch from a certain student *i* and *X*_*j*_, an overlapping epoch from another student *j*, TI value was calculated as follows.

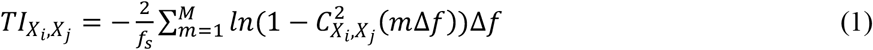

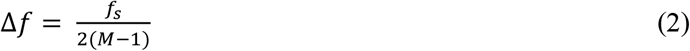

Here, *C_X_*_*i*_,*X*_*j*_ () is the magnitude squared coherence calculation, *f*_*s*_ is the sampling rate, M is the number of desired frequency points in the interval between 0 and the Nyquist frequency 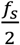. The Δ*f* is the frequency resolution. Theta, alpha, and low-beta and high-beta TI were computed by summing the coherence values within 4-8 Hz, 8-13 Hz, 13-18 Hz, and 18-30 Hz, respectively.

TI for one pair of students for each session was obtained by averaging TI values across all epochs and the two recording electrodes. A minimum of 6 artifact-free common epochs for paired students were included for further analysis. The lower limit was empirically chosen to get a comparable minimum data amount for each pair of students with the previous studies^4,16^. 96.6% of TI values for each pair and each epoch remained for the following analysis. A *N* * *N* pairwise TI matrix (*N* is the number of students) could be obtained for each session. TI values within the matrix were then normalized to [0,1] for each session, following the practice in previous studies ^4,16^. Then, the matrixes were averaged across *K* sessions to obtain an averaged inter-brain coupling for each pair of students for a specific discipline (*K* = 18 for Chinese and *K* = 20 for Math).

Then, student-class coupling for student *i* was obtained by averaging TI values over all possible pairwise combinations between the student *i* and the rest of the class. Student-excellence coupling for student *i* was computed by averaging TI values over all possible pairwise combinations between the student *i* and the excellences except themselves if included. Therefore, for each student, there would be a student-class coupling value and a student-excellence coupling value as indicators of disciplinary learning process for the whole semester for each frequency band.

Furthermore, Pearson’s correlations between individuals’ student-class coupling (or student-excellence coupling) and their corresponding learning outcomes were calculated for soft and hard disciplines separately. For student-class coupling, all students were included in the correlation analysis. For student-excellence coupling, however, the excellences themselves were not included during the correlation analysis between student-excellence coupling and learning outcomes. For example, if individuals’ student-excellence coupling was computed with the top *N*_*E*_ students, then the top *N*_*E*_ students’ coupling values, as well as their learning outcomes, would be removed, leaving (*N* − *N*_*E*_) out of the *N* students for conducting correlations. The effect of excellence number on the relationship between student-excellence coupling and learning outcomes was analyzed.

Additionally, we conducted similar correlational analyses between single-brain EEG features and the final exam scores for comparison. The relative power of each frequency band of interest (theta, alpha, and low-beta and high-beta) was obtained by dividing the power in the 1-40 Hz band after a fast Fourier transform for each 30-second epoch. Then, values of the relative power of each frequency band of interest were averaged across all epochs and all sessions within each discipline for each student as the single-brain EEG features. Finally, Pearson’s correlations between individuals’ single-brain EEG features and corresponding learning outcomes were calculated separately for soft and hard disciplines.

## Acknowledgement

We would like to thank Professor Yongdi Zhou and Zhuoran Li for their comments on the manuscript. This work was supported by the National Natural Science Foundation of China (61977041, 62107025 and 62177030), and Tsinghua University Spring Breeze Fund (2021Z99CFY037).

## Data availability

The raw data will be available upon reasonable request to the corresponding author due to underage privacy protection. The minimum de-identified dataset used to generate the findings of this study are available at : https://cloud.tsinghua.edu.cn/d/83a776b5db1349c29fd7/.

## Code availability

The Matlab code for analysis and figures generation are available at: https://cloud.tsinghua.edu.cn/d/83a776b5db1349c29fd7/.

## Author contributions

D.Z. and Y.Z. designed the experiment and revised the manuscript. J.C. analyzed the data and wrote the manuscript. Q.P pre-analzed the data. X.G. and B.L. collected the data.

## Supplementary Materials

**Fig.S1:**
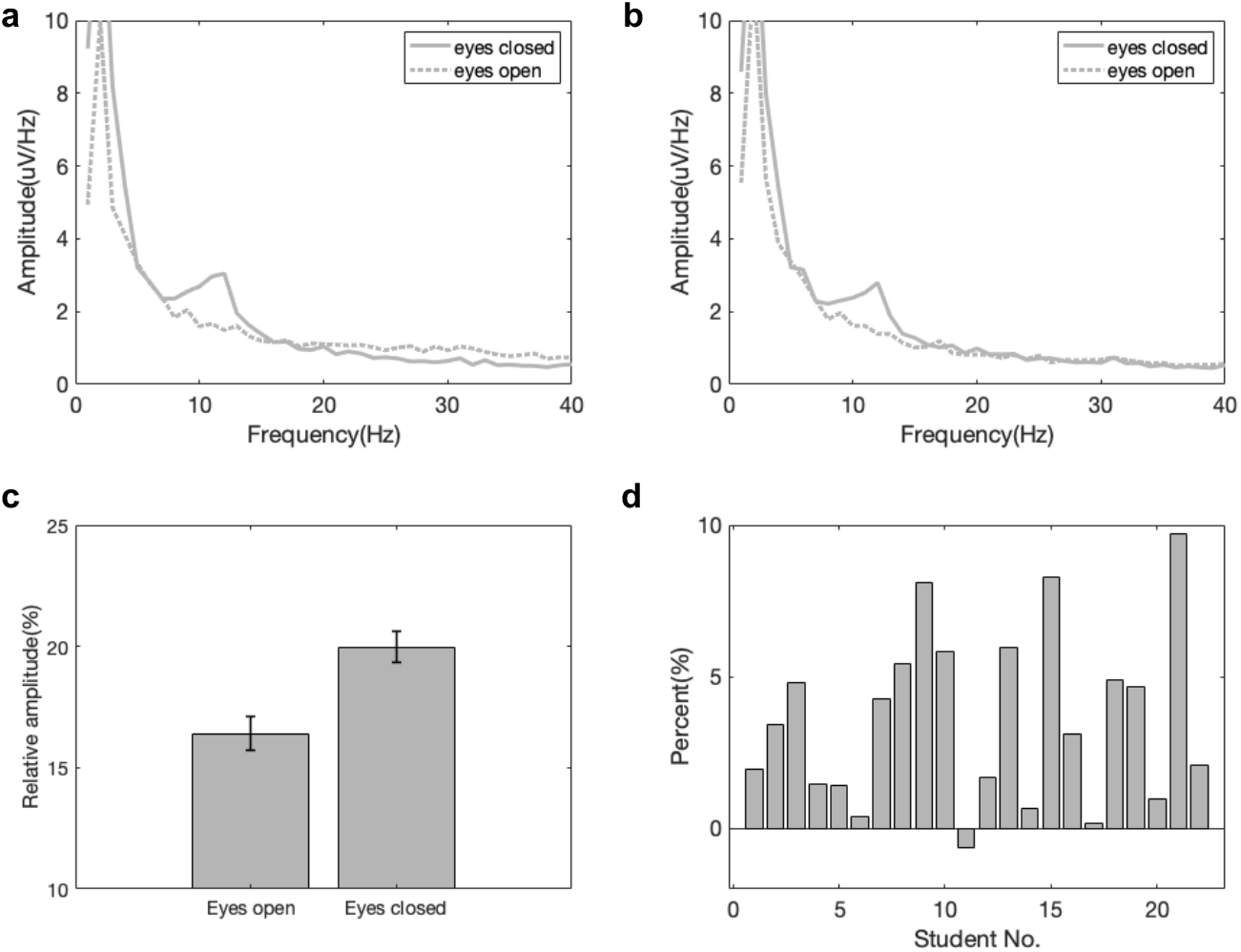
The signal quality validation of EEG headbands in a 2-minute eye-closed/open task. Twenty-two students out of the same class volunteered to participate in this task. Students were required to open and close eyes for two minutes respectively when sitting in their classroom. Then, fast Fourier transform was conducted to compare the frequency spectral characteristics of EEG signals between conditions. **a**, The frequency spectra for a representative student at Fp1. The solid line represents the eye-closed condition, and the dashed lines represent the eye-open condition. **b**, The frequency spectra for the same student at Fp2. **c**, Relative amplitudes of the alpha band (8-13 Hz) in the eye-open and eye-closed conditions for all the students. Errorbar indicates standard deviation. The amplitudes were averaged across Fp1 and Fp2; **d**, Percent of relative amplitude differences (eye-closed minus eye-open) at the alpha band for each student. The amplitudes were averaged across Fp1 and Fp2.

**Fig.S2.**
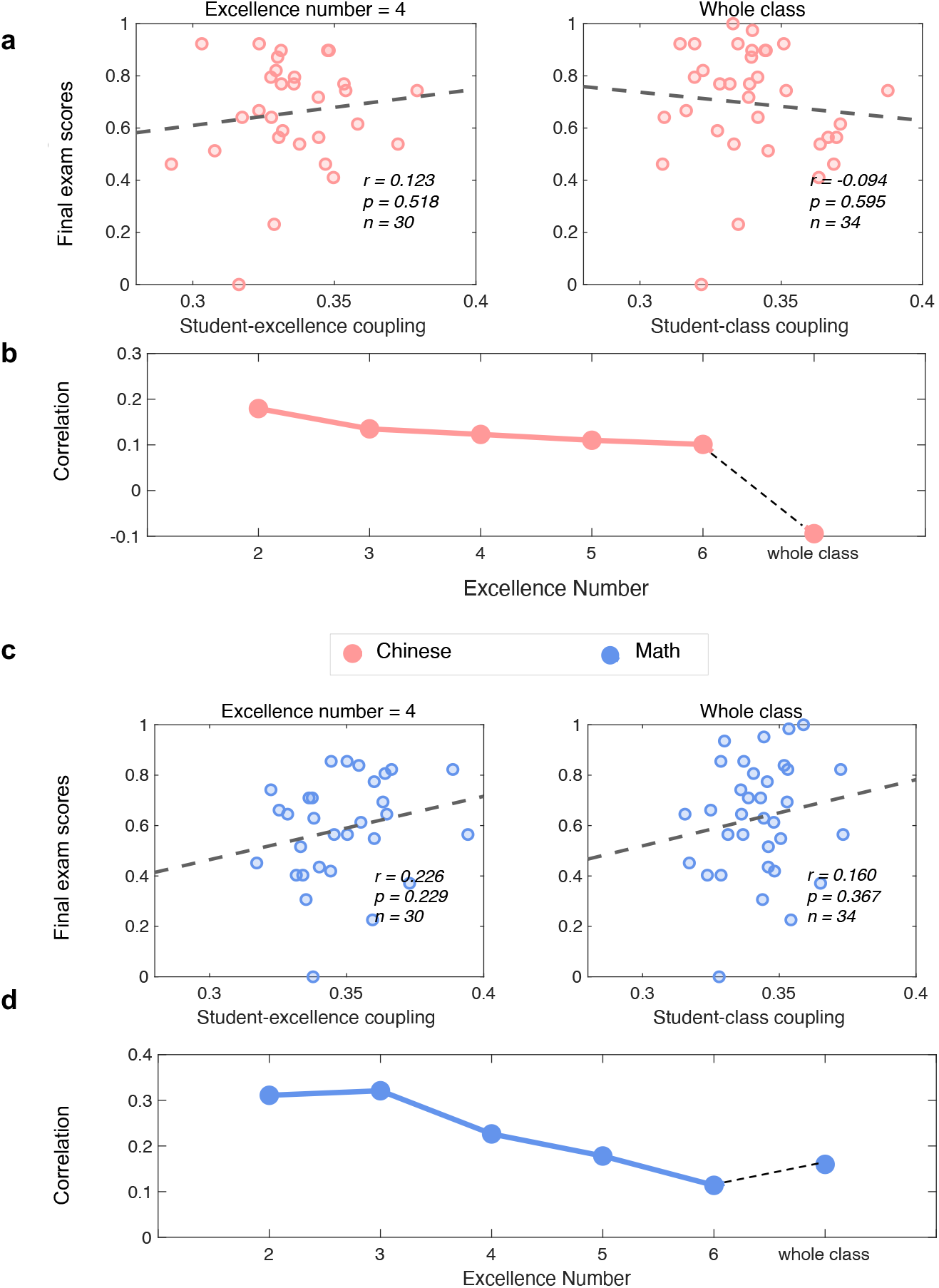
The correlations between student-excellence/class coupling with one’s final exam score of Chinese and Math at 1-20 Hz. **a**, Scatter plots between student-excellence coupling (left, excellence number = 4) and student-class coupling (right) and the final exam score of Chinese at 1-20 Hz. **b**, Correlation r values as a function of the number of excellences included in the calculation of student-excellence coupling. **c**, Scatter plots between student-excellence coupling (left, excellence number = 4) and student-class coupling (right) and the final exam scores of Math at 1-20 Hz. **d**, Correlation r values as a function of the number of excellences included in the calculation of student-excellence coupling. Note that the excellences themselves were not included in the correlation analysis.

**Fig.S3:**
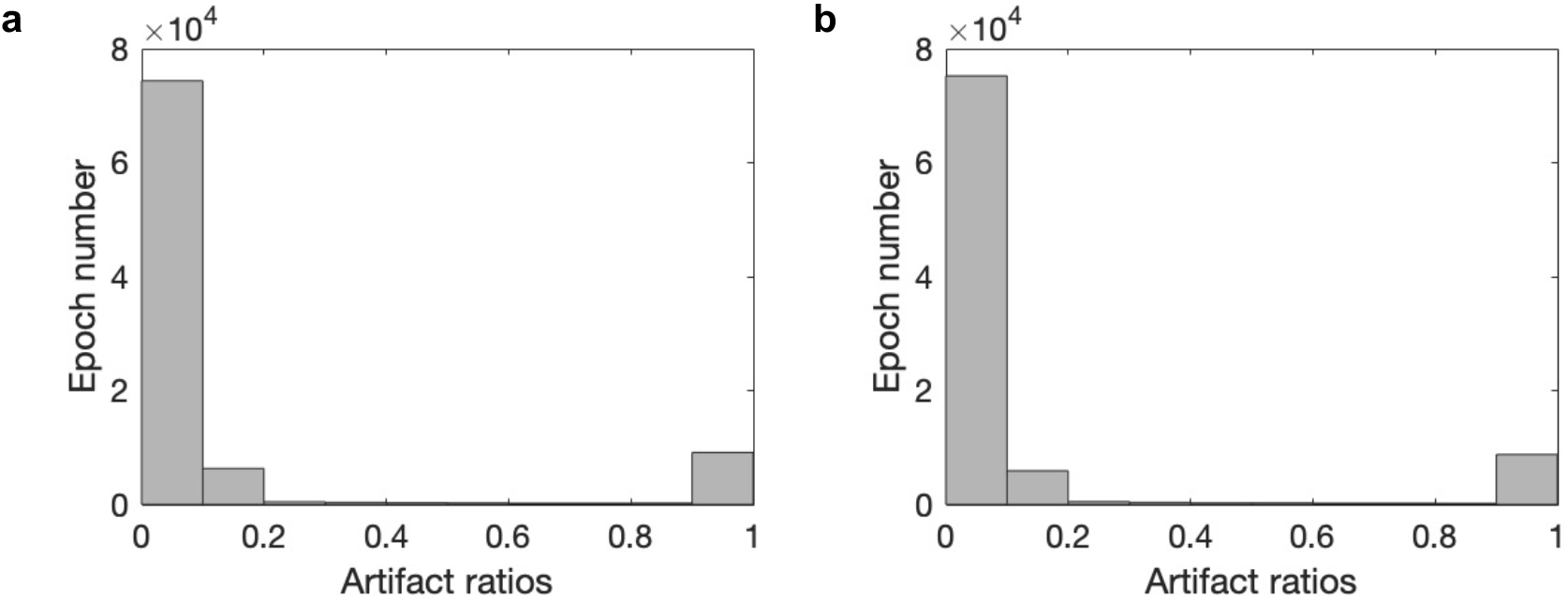
The distribution of the artifacts ratios per epoch for (**a**) Fp1 and (**b**) Fp2. The ratios for saturated samples per epoch illustrated a 2-tailed distribution.

**Fig.S4:**
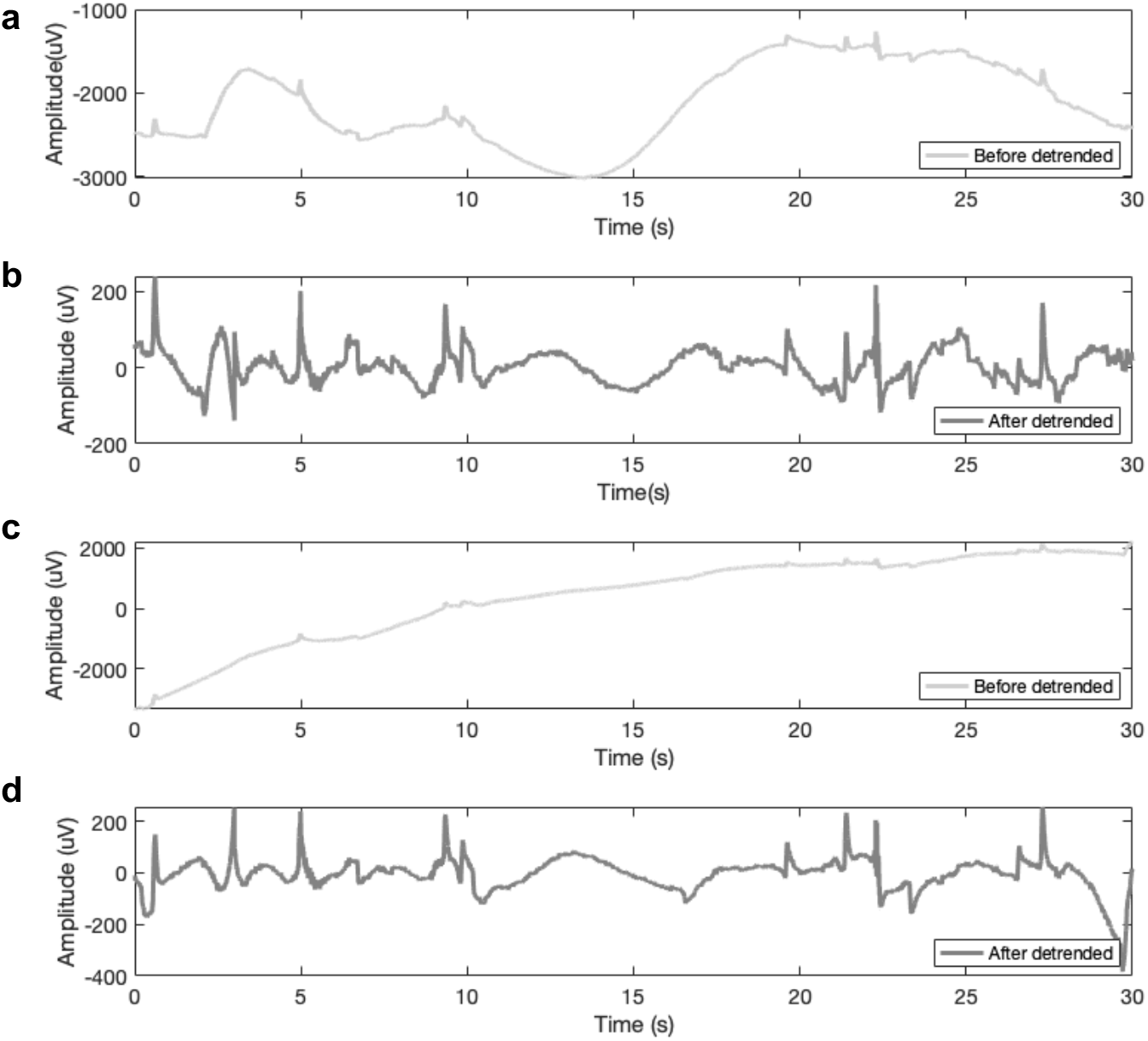
An illustration of a representative EEG epoch before and after the detrending procedure with the Noisetool toolbox (**a, b**) at Fp1; (**c, d**) at Fp2. The lines with a lighter color indicated the condition before detrending. The lines with a darker color indicated the condition after detrending. The slow drift was removed after the detrending procedure.

**Fig.S5:**
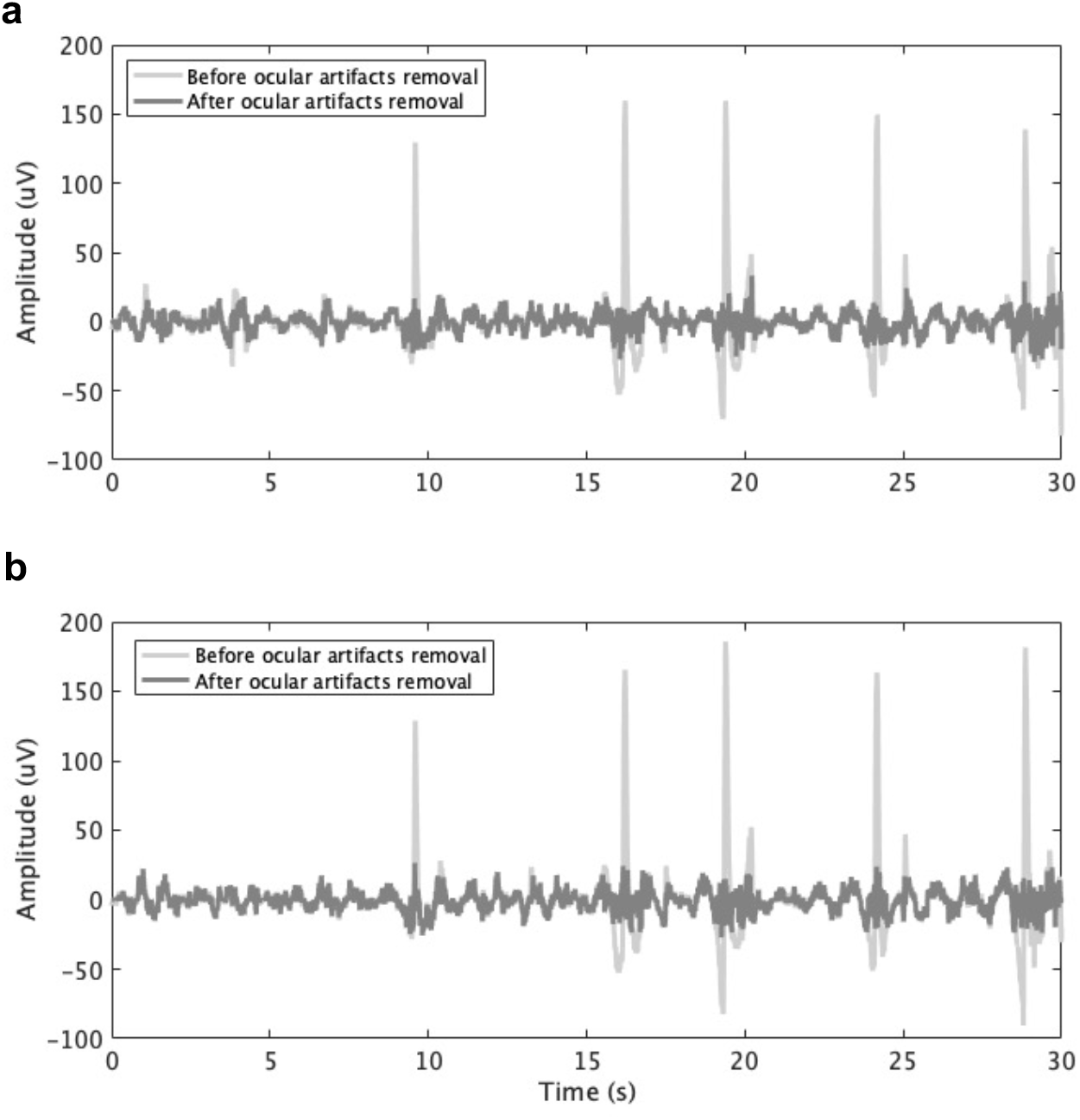
An illustration of a representative EEG epoch before and after the ocular artifact removal procedure with the MSDL toolbox (**a**) at Fp1; (**b**) at Fp2. The lines with a lighter color indicated the condition before ocular artifact removal. The lines with a darker color indicated the condition after ocular artifact removal. The ocular artifact was removed after the procedure.

**Fig.S6:**
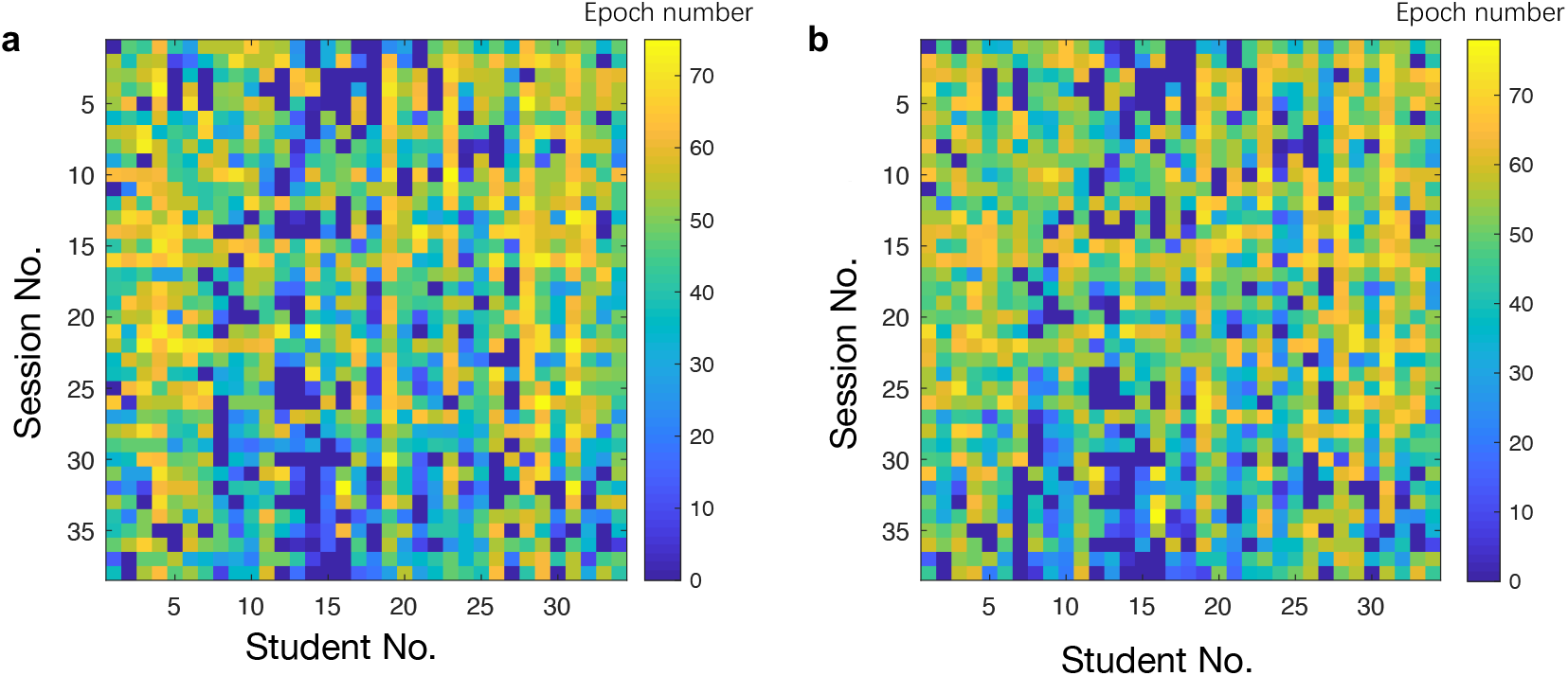
The number of retained epochs per session per student (**a**) at Fp1; (**b**) at Fp2 after the preprocessing procedure. The colorbar indicated the number of retained epochs.

## Notes

### Competing Interest Statement

The authors have declared no competing interest.

### Summary of Updates

we revised the introduction and discussions

## Reference

1. Darling-Hammond, L., Flook, L., Cook-Harvey, C., Barron, B. & Osher, D. Implications for educational practice of the science of learning and development. Applied Developmental Science 24, 97–140 (2020).

2. Valiente, C., Swanson, J., DeLay, D., Fraser, A. M. & Parker, J. H. Emotion-related socialization in the classroom: Considering the roles of teachers, peers, and the classroom context. Developmental psychology 56, 578 (2020).

3. Vandenbroucke, L., Spilt, J., Verschueren, K., Piccinin, C. & Baeyens, D. The classroom as a developmental context for cognitive development: A meta-analysis on the importance of teacher–student interactions for children’s executive functions. Review of Educational Research 88, 125–164 (2018).

4. Dikker, S. et al. Brain-to-brain synchrony tracks real-world dynamic group interactions in the classroom. Current biology 27, 1375–1380 (2017).

5. Biglan, A. Relationships between subject matter characteristics and the structure and output of university departments. Journal of applied psychology 57, 204 (1973).

6. Lindblom-Ylänne, S., Trigwell, K., Nevgi, A. & Ashwin, P. How approaches to teaching are affected by discipline and teaching context. Studies in Higher education 31, 285–298 (2006).

7. Neumann, R., Parry, S. & Becher, T. Teaching and learning in their disciplinary contexts: A conceptual analysis. Studies in higher education 27, 405–417 (2002).

8. Rosman, T., Mayer, A.-K., Kerwer, M. & Krampen, G. The differential development of epistemic beliefs in psychology and computer science students: A four-wave longitudinal study. Learning and Instruction 49, 166–177 (2017).

9. Smith*, S. N. & Miller, R. J. Learning approaches: Examination type, discipline of study, and gender. Educational psychology 25, 43–53 (2005).

10. Hofer, B. K. Dimensionality and disciplinary differences in personal epistemology. Contemporary educational psychology 25, 378–405 (2000).

11. Arbaugh, J. B. Does academic discipline moderate CoI-course outcomes relationships in online MBA courses? The Internet and Higher Education 17, 16–28 (2013).

12. Foung, D. & Chen, J. Discovering disciplinary differences: blending data sources to explore the student online behaviors in a University English course. Information Discovery and Delivery (2019).

13. Cohen, S. S. et al. Neural engagement with online educational videos predicts learning performance for individual students. Neurobiology of learning and memory 155, 60–64 (2018).

14. Meshulam, M. et al. Neural alignment predicts learning outcomes in students taking an introduction to computer science course. Nature communications 12, 1–14 (2021).

15. Davidesco, I. et al. Brain-to-brain synchrony predicts long-term memory retention more accurately than individual brain measures. BioRxiv 644047 (2019).

16. Bevilacqua, D. et al. Brain-to-brain synchrony and learning outcomes vary by student–teacher dynamics: Evidence from a real-world classroom electroencephalography study. Journal of cognitive neuroscience 31, 401–411 (2019).

17. Brandt, N. D., Lechner, C. M., Tetzner, J. & Rammstedt, B. Personality, cognitive ability, and academic performance: Differential associations across school subjects and school tracks. Journal of personality 88, 249–265 (2020).

18. Marton, F. & Säljö, R. On qualitative differences in learning: I—Outcome and process. British journal of educational psychology 46, 4–11 (1976).

19. Shamay-Tsoory, S. G. Brains that fire together wire together: interbrain plasticity underlies learning in social interactions. The Neuroscientist 1073858421996682 (2021).

20. Shamay-Tsoory, S. G. & Mendelsohn, A. Real-life neuroscience: an ecological approach to brain and behavior research. Perspectives on Psychological Science 14, 841–859 (2019).

21. Sonkusare, S., Breakspear, M. & Guo, C. Naturalistic stimuli in neuroscience: critically acclaimed. Trends in cognitive sciences 23, 699–714 (2019).

22. De Sanctis, P. et al. Time to move: Brain dynamics underlying natural action and cognition. European Journal of Neuroscience (2021).

23. Xu, J. & Zhong, B. Review on portable EEG technology in educational research. Computers in Human Behavior 81, 340–349 (2018).

24. Janssen, T. W. et al. Opportunities and limitations of mobile neuroimaging technologies in educational neuroscience. Mind, Brain, and Education (2021).

25. Davidesco, I. Brain-to-brain synchrony in the STEM classroom. CBE—Life Sciences Education 19, es8 (2020).

26. Ko, L.-W., Komarov, O., Hairston, W. D., Jung, T.-P. & Lin, C.-T. Sustained attention in real classroom settings: An EEG study. Frontiers in human neuroscience 11, 388 (2017).

27. Babiker, A., Faye, I., Mumtaz, W., Malik, A. S. & Sato, H. EEG in classroom: EMD features to detect situational interest of students during learning. Multimedia Tools and Applications 78, 16261–16281 (2019).

28. Davidesco, I., Matuk, C., Bevilacqua, D., Poeppel, D. & Dikker, S. Neuroscience Research in the Classroom: Portable Brain Technologies in Education Research. Educational Researcher 50, 649–656 (2021).

29. Jang, K.-I., Lee, S., Lee, S.-H. & Chae, J.-H. Frontal alpha asymmetry, heart rate variability, and positive resources in bereaved family members with suicidal ideation after the Sewol ferry disaster. Psychiatry investigation 15, 1168 (2018).

30. Kang, J.-S., Ojha, A. & Lee, M. Development of Intelligent Learning Tool for Improving Foreign Language Skills Based on EEG and Eye Tracker. in Proceedings of the 3rd international conference on human-agent interaction 121–126 (2015).

31. Kang, J.-S., Ojha, A. & Lee, M. Concentration monitoring with high accuracy but low cost EEG device. in International Conference on Neural Information Processing 54–60 (Springer, 2015).

32. Kwon, J.-W. et al. Intraoperative real-time stress in degenerative lumbar spine surgery: simultaneous analysis of electroencephalography signals and heart rate variability: a pilot study. The Spine Journal 20, 1203–1210 (2020).

33. Kwon, J.-W. et al. Which Factors Affect the Stress of Intraoperative Orthopedic Surgeons by Using Electroencephalography Signals and Heart Rate Variability? Sensors 21, 4016 (2021).

34. Wen, X., Mo, J. & Ding, M. Exploring resting-state functional connectivity with total interdependence. Neuroimage 60, 1587–1595 (2012).

35. Chabin, T. et al. Interbrain emotional connection during music performances is driven by physical proximity and individual traits. Annals of the New York Academy of Sciences (2021).

36. Cavanagh, J. F. & Frank, M. J. Frontal theta as a mechanism for cognitive control. Trends in cognitive sciences 18, 414–421 (2014).

37. Cavanagh, J. F. & Shackman, A. J. Frontal midline theta reflects anxiety and cognitive control: meta-analytic evidence. Journal of physiology-Paris 109, 3–15 (2015).

38. Eschmann, K. C., Bader, R. & Mecklinger, A. Topographical differences of frontal-midline theta activity reflect functional differences in cognitive control abilities. Brain and cognition 123, 57–64 (2018).

39. Clayton, M. S., Yeung, N. & Kadosh, R. C. The roles of cortical oscillations in sustained attention. Trends in cognitive sciences 19, 188–195 (2015).

40. Jensen, O. & Tesche, C. D. Frontal theta activity in humans increases with memory load in a working memory task. European journal of Neuroscience 15, 1395–1399 (2002).

41. Maurer, U. et al. Frontal midline theta reflects individual task performance in a working memory task. Brain topography 28, 127–134 (2015).

42. Sauseng, P., Griesmayr, B., Freunberger, R. & Klimesch, W. Control mechanisms in working memory: a possible function of EEG theta oscillations. Neuroscience & Biobehavioral Reviews 34, 1015–1022 (2010).

43. Williams, C. C., Kappen, M., Hassall, C. D., Wright, B. & Krigolson, O. E. Thinking theta and alpha: Mechanisms of intuitive and analytical reasoning. NeuroImage 189, 574–580 (2019).

44. Haag, L. & Götz, T. Mathe ist schwierig und Deutsch aktuell: Vergleichende Studie zur Charakterisierung von Schulfächern aus Schülersicht. Psychologie in Erziehung und Unterricht 59, 32–46 (2012).

45. Sauseng, P. et al. EEG alpha synchronization and functional coupling during top-down processing in a working memory task. Human brain mapping 26, 148–155 (2005).

46. Fink, A. & Benedek, M. EEG alpha power and creative ideation. Neuroscience & Biobehavioral Reviews 44, 111–123 (2014).

47. Lustenberger, C., Boyle, M. R., Foulser, A. A., Mellin, J. M. & Fröhlich, F. Functional role of frontal alpha oscillations in creativity. Cortex 67, 74–82 (2015).

48. Cooper, N. R., Burgess, A. P., Croft, R. J. & Gruzelier, J. H. Investigating evoked and induced electroencephalogram activity in task-related alpha power increases during an internally directed attention task. Neuroreport 17, 205–208 (2006).

49. Adams, W. K. & Wieman, C. E. Development and validation of instruments to measure learning of expert-like thinking. International Journal of Science Education 33, 1289–1312 (2011).

50. Szulewski, A. et al. Starting to think like an expert: an analysis of resident cognitive processes during simulation-based resuscitation examinations. Annals of Emergency Medicine 74, 647–659 (2019).

51. Petrilli, M. J. All together now? Education high and low achievers in the same classroom. Education Next 11, 48–56 (2011).

52. VanTassel-Baska, J. & Stambaugh, T. Challenges and possibilities for serving gifted learners in the regular classroom. Theory into practice 44, 211–217 (2005).

53. Westberg, K. L. & Daoust, M. E. The results of the replication of the classroom practices survey replication in two states. The National Research Center on the Gifted and Talented Newsletter 3, (2003).

54. Glass, T. F. What gift?: The reality of the student who is gifted and talented in public school classrooms. Gifted Child Today 27, 25–29 (2004).

55. Pan, Y., Cheng, X. & Hu, Y. Three heads are better than one: Cooperative learning brains wire together when a consensus is reached. bioRxiv (2021).

56. Wang, L., Li, M., Yang, T. & Zhou, X. Mathematics Meets Science in the Brain. Cerebral Cortex (2021).

57. de Cheveigné, A. & Arzounian, D. Robust detrending, rereferencing, outlier detection, and inpainting for multichannel data. NeuroImage 172, 903–912 (2018).

58. Kanoga, S., Kanemura, A. & Asoh, H. Multi-scale dictionary learning for ocular artifact reduction from single-channel electroencephalograms. Neurocomputing 347, 240–250 (2019).

